# *G-protein coupled receptor 88* knock-down in the associative striatum reduces the psychiatric symptoms in a translational model of Parkinson’s disease

**DOI:** 10.1101/617456

**Authors:** Benjamin Galet, Manuela Ingallinesi, Jonathan Pegon, Anh Do Thi, Philippe Ravassard, Nicole Faucon Biguet, Rolando Meloni

## Abstract

Beyond the motor disability, Parkinson’s disease (PD) is also characterized by an early appearance of psychiatric symptoms such as apathy, depression, anxiety and cognitive deficits, which can entail dementia and psychosis in later stages. While current treatments may provide some level of symptomatic relief, their use is limited by the development of adverse effects such as impulse-control disorders. There is thus a medical need for targets with novel modes of action to treat these aspects of PD. In this context, we investigated GPR88, an orphan G-protein coupled receptor that is associated with psychiatric disorders and highly enriched in the striatum, where it exerts an inhibitory control over neurotransmitter systems that are compromised in PD. To evaluate the potential of GPR88 as a target for the treatment of the psychiatric symptoms of PD, we knocked-down (KD) its expression in sensorimotor (dorsolateral, DLS) or associative (dorsomedial, DMS) striatal areas in a translational rat model of early PD. Our findings indicate that *Gpr88-*KD in the DMS, but not DLS, reduced the alterations in mood, motivation and cognition that characterized the model, through modulation of the expression of *regulator of G-protein signaling 4* (*Rgs4*) and of transcription factor ΔFosB. Furthermore, the rat model of PD exhibited allostatic changes in striatal activity markers that may be related to patterns observed in patients, and which were reduced by *Gpr88-*KD. Taken together, these results thus highlight the relevance of GPR88 as a therapeutic target for the psychiatric symptoms of PD.

## INTRODUCTION

While Parkinson’s disease (PD) is classically defined as a motor disorder, it also entails psychiatric symptoms that are a major burden on patients’ quality of life (1). Many of these symptoms have been grouped as part of a “hypodopaminergic” syndrome resulting from the loss of nigrostriatal neurons, which includes apathy, depression, anxiety and cognitive impairment, and affects 35 to 85% of patients depending on the symptom (2, 3). Apathy in particular, defined as a decrease in motivational drive, has emerged as a core symptom of PD, and is present in up to 70% of patients (4). It is often associated with cognitive impairment, as apathetic patients are found to have worse executive dysfunction and an increased risk of developing dementia in later stages (5). Psychosis also develops in up to 60% of patients, and is an additional risk factor for dementia(6).

A remarkable feature of these symptoms is that they frequently emerge years before the onset of motor impairments (7, 8). At the time of diagnosis, studies have for instance found that 37% of patients already suffered from depression, 27% from apathy, 17% from anxiety (9), 20-40% from cognitive dysfunction (10), and 42% reported minor psychotic phenomena, such as passage and presence hallucinations (11).

While dopaminergic medications can efficiently treat some of these aspects (3, 4), their effect is hindered by the frequent development of “hyperdopaminergic” symptoms such as impulse-control disorders, which can have disastrous consequences (2). Furthermore, existing treatments for cognitive impairment and psychosis have variable efficacy and limiting side effects (6, 12). There is thus a crucial unmet medical need for the development of new symptomatic treatments to manage these symptoms.

In this context, the orphan G-protein Coupled Receptor 88 (GPR88) is emerging as a particularly suited target. *GPR88* has been associated with Bipolar Disorder and Schizophrenia (13, 14), and deleterious mutations were found to induce profound learning deficits and a hyperkinetic movement disorder in humans (15). GPR88 is mainly expressed in striatal Medium Spiny Neurons (MSN) at the level of the corticostriatal synapse (16), where it exerts an inhibitory control over monoamine and neuropeptide neurotransmission through G_i/o_ coupling (17). Indeed, *Gpr88*-KO mice are hypersensitive to D2 agonists, and their MSNs exhibit increased signaling of delta opioid and muscarinic receptors (18, 19). Given these characteristics, it is not surprising that *Gpr88*-KO mice were found to have impaired affective, goal-directed, cognitive and sensorimotor behaviors (18–22).

GPR88’s restricted localization and wide-ranging effects have thus made it an interesting therapeutic target for disorders of the basal ganglia. For instance, preclinical studies have already revealed promising results in a model of schizophrenia (23) and potential involvement in alcohol use disorder (24). Given that GPR88 exerts inhibitory control over the key neurotransmitter systems that are altered in PD, we hypothesized that inhibiting GPR88 in the striatum could have a therapeutic effect by potentiating endogenous neurotransmission.

In order to evaluate GPR88’s potential as a therapeutic target for the psychiatric symptoms of PD, we first developed a translational model of early PD in rats by reproducing the loss of dopamine (DA) that is observed in the first stages of the disease, specifically affecting sensorimotor territories of the striatum (25), through stereotaxic injections of the 6-hydroxydopamine (6OHDA) toxin. As no antagonists are currently available for this receptor, we then used a localized gene therapy approach to knock-down the expression of *Gpr88* (*Gpr88-*KD). This strategy enabled us to assess the effects of *Gpr88-*KD in the sensorimotor and associative areas of the striatum independently (dorsolateral and dorsomedial striatum in rodents, respectively), the latter having been frequently associated with psychiatric symptoms in patients (26). The effects of both procedures were then evaluated at behavioral and molecular levels.

## MATERIALS AND METHODS

### Animals

Animal studies were authorized by an ethical committee reporting to the French Ministry of Agriculture (APAFIS reference #3669-2016011817516297) and were conducted in the in-house SPF animal facility, which was approved by the Veterinary Inspection Office (agreement reference B75-13-19). The animals were handled throughout the study in compliance with the European Union 2010 Animal Welfare Act, and the 2010/63 French directive. The procedures were reported following the ARRIVE guidelines (27).

The experiments were performed with male Sprague-Dawley rats that weighed 300g on average at the time of the first surgery (7 weeks old). The rats were first housed in groups of four until the first surgery, after which they were housed in pairs. Animal well-being was checked daily by the experimenters or the animal facility staff. The cages were ventilated and enriched with cardboard tunnels, and food and water were available *ad libitum*. The facility used a 12h day/night cycle starting at 8 AM, and controlled the temperature and the humidity levels daily.

### Study design

A schematic representation of the study design and timeline is presented in the supplementary figures (**Fig. S1**). Two hypotheses were emitted at the beginning of the project, namely that (i) the partial lesion of the rat DLS would reproduce some of the psychiatric symptoms of PD, and that (ii) inactivating *Gpr88* in different striatal areas of the lesioned rats would affect different behavioral parameters. In order to test each hypothesis, several experimental groups were designed. (i) To assess the effect of the loss of DA, two groups of animals were compared: 6OHDA-injected *vs* SHAM-injected. The rats from both groups were also transduced with an inactive control LV miR-neg sequence in the different striatal compartments. (ii) To determine the effects of *Gpr88* inactivation in 6OHDA-lesioned rats, stereotaxic injections of lentiviruses were performed either in the dorsolateral (DLS), dorsomedial (DMS) or ventral striatum (VS; Nucleus Accumbens Core). However, as experiments were underway, no major effects of *Gpr88* inactivation in the ventral striatum were observed, while intriguing interactions were emerging between the dorsolateral and dorsomedial tiers of the striatum. For the sake of clarity, we thus decided to limit the scope of this article to the DLS-DMS interactions in both the effects of the lesion and of *Gpr88-*KD.

The data were accumulated over 17 replication batches. Each batch consisted of 6-10 animals that were randomly distributed across the different experimental conditions, making sure that no 2 rats of the same conditions were housed together. In order to comply with the 3Rs guidelines, the appropriate sample size was calculated based on behavioral data from preliminary experiments, using the G*Power3 software (28). To reach a statistical power of 0.9 with the alpha level set at 0.05, the recommended sample size was of 9-10 animals per condition, depending on the behavioral task. Replication batches were thus stopped once the recommendation was met.

Finally, we observed an important level of cumulative variability in the quality and extent of the lesions and lentiviral transductions. In order to limit the effect of this variability and enhance the reproducibility of our findings, strict inclusion criteria were applied. First of all, the DA loss had to affect at least 20% of the DLS, but no more than 15% of the neighboring DMS and VS. Next, sufficient GFP fluorescence had to be present in the area targeted with LV miR-*Gpr88*. Furthermore, in the rare occurrences of adverse events (such as important weight loss or inflammatory reactions), the rats were excluded from the statistical analyses.

### Stereotaxic injections of 6OHDA and lentiviral vectors

Before the beginning of the surgeries, the animals were first anaesthetized with 4% isoflurane (IsoVet, Osalia) in an induction chamber (Minerve Veterinary Equipment, Esternay, France) for 5 minutes before being placed into a Kopf stereotaxic surgery apparatus (Phymep, Paris, France). Anesthesia was maintained throughout the surgery with an isoflurane pump (Univentor, Zejtun, Malta). General and localized analgesia was induced with subcutaneous injections of Buprecare (buprenorphine; 0,05mg/kg) (Axience) and Xylovet (lidocaine; 17,5mg/kg) (Ceva) before beginning the surgery. 4µl of a solution of 6OHDA (3µg/µl, 12µg total per hemisphere) in saline + 0.02% ascorbic acid (all chemicals from Sigma) or of a control solution (saline + 0.02% ascorbic acid only) were then injected bilaterally in the DLS using a 10µL Hamilton syringe (Phymep, Paris, France). The coordinates were as follow: anteroposterior (AP) + 0,7mm; mediolateral (ML) ± 3,8mm; dorsoventral (DV) −5,5mm (2µl) and −4,5mm (remaining 2µl) from bregma(29). The injections were performed at a rate of 0.5µl/min, and the syringe was left in place for 3 minutes after the end of each injection to allow for proper diffusion of the toxin. Following the surgeries, the wellbeing of the animals was checked daily. While some rats transiently lost weight, they typically recovered within 3 to 5 days.

### Lentivirus production and stereotaxic injection

The lentiviral vectors were generated at the in-house iVector platform, using the BLOCK-iT Pol II miR RNAi expression vector kit (Invitrogen). The vector construct contained an engineered miR sequence to drive *Gpr88* knock-down. The modified miR consisted of a shRNA inserted within the miRNA 155 flanking sequences. One of the shRNA complementary sequences was designed to target either *Gpr88* mRNA (“miR-*Gpr88*”), or a control sequence that is not expressed in the genome (“miR-neg”). The use of a shRNA sequence allowed for high specificity of RNA interference under control of the PGK promoter, which is best suited for *in vivo* studies. The lentiviruses were stored in PBS at −80°C, at an average of 1,6×10^5^ transducing units (TU) /µL.

6µL of the lentivirus solution were bilaterally injected in the DLS or DMS two weeks after the 6OHDA injections, following the same general surgical procedure. The coordinates were however different, as the lentiviruses were injected at 4 different sub-sites per hemisphere (8 total) to insure sufficient knock-down of *Gpr88* expression. When targeting the DLS, the coordinates were the following: (1) AP +1,2 mm; ML ±3,6 mm; DV −5,5mm and −4,5mm (2) AP +0,2 mm, ML ±4mm, DV −5,5mm and −4,5mm from bregma. Regarding the DMS, the coordinates were: (1) AP +1,2 mm; ML ±2 mm; DV −5,5mm and −4,5mm (2) AP +0,2 mm, ML ±2,2mm, DV −5,5mm and −4,5mm from bregma.

### Behavioral tests

Each test was set up by the same experimenter and around the same time of day for each batch of animals, in between 9 AM and 4 PM. The experimenter was unaware of the status of the rats, which were brought in the testing rooms one hour before the beginning of the task, in order for them to acclimate to the environment. The luminosity was controlled for every experiment (35 lux), and the testing apparatus were cleaned before each test and between rats with a disinfectant solution (Aniospray, Dutscher).

#### Actimeter

General motor and exploratory behavior was assessed for 15 minutes using the Panlab Infrared Actimeter (Harvard Apparatus, Holliston, MA, USA). Horizontal, stereotyped and vertical movements were automatically quantified and cumulated into 5-minute segments.

#### Sucrose preference

Rats were isolated in enriched individual cages for 72h, during which time they were given access to two bottles containing either tap water, or tap water supplemented with 0,5% sucrose (Sigma). The first 24h were considered as an acclimation phase, and were not included in the analysis. After 48h, the position of the bottles was inverted to avoid side preference effects. Bottles were weighed daily in order to calculate the amount of consumed liquids. Sucrose preference was calculated as the percentage of sucrose intake / total intake. However, as the level of sucrose preference exhibited by control rats was lower than what we had observed during pilot experiments (at 0,25% sucrose), the concentration had to be re-evaluated after several batches of animals (0,5%). For this reason, the number of data points for this experiment is relatively lower. General consummatory behavior was also tracked by weighing the food dispenser at the beginning and end of the isolation phase.

#### Social Novelty Discrimination (SND)

Social interaction and selective attention were then evaluated using the social novelty discrimination task (SND) as previously described (23, 30). As this behavioral paradigm requires a preliminary isolation phase, we performed it at the end of the sucrose preference test. Briefly, a first juvenile was placed into the home cage of the tested rat for a presentation period (P1) of 30 min. The time spent by the tested rat investigating the juvenile (anogenital sniffing, pursuing, allogrooming) was timed manually for the first 5 min. At the end of P1, a second juvenile was introduced in the cage, and the time spent investigating the novel vs the familiar juvenile was timed by the experimenter (presentation period P2). The “discrimination ratio” was calculated as the time spent by the tested rat interacting with the novel juvenile over the familiar juvenile during P2.

#### Prepulse Inhibition (PPI)

Sensorimotor gating was assessed using an auditory prepulse inibition apparatus (IMETRONIC, Pessac, France). The adapted protocol (31) contained three phases: a first acclimation period of 10 minutes, followed by a phase of habituation to the startle stimulus, and ending with the testing phase. During the acclimation phase, a background white noise of 60dB was played, which persisted throughout the whole testing session. The habituation phase consisted of 10 startle-inducing auditory “pulses” played at 110dB (7Khz, 100ms), and at random intervals between 15-30 seconds. During the testing phase, four different types of stimuli were presented: a pulse alone (110dB), a prepulse-pulse pairing, a prepulse alone, or no sound (to assess background movement). The prepulses (20ms, 70-80dB) preceded the startle pulses by 100ms. Each condition was presented 10 times in a pseudo-randomized order, at random intervals between 15 and 30 seconds. Prepulse inhibition (%PPI) was measured as the reduction in startle response during prepulse–pulse conditions compared to pulse-alone trials.

#### Forced Swim Test (FST)

The rats were placed in a transparent cylinder filled up to 35 cm with 24°C (±1) water for 5 minutes, and recorded using a digital camera. A trained experimenter blinded to the conditions then analyzed the behavior of the animals using a previously described sampling method (32), calculating immobility, swimming, climbing and diving counts.

### Immunolabelling and in situ hybridization

Following the end of the behavioral procedures (after 2-4 days), the rats were anaesthetized in an isoflurane chamber and decapitated. The brains were rapidly removed and snap-frozen for 90 seconds in isopentane at −55°C (Carlo Erba Reagents). Coronal cryosections (12µm) of the striatum were used to control for the presence and extent of GFP signal in the targeted areas. The slices were then post-fixed in 4% PFA for 30 minutes at 4°C in order to perform immunolabelling and *in situ* hybridization experiments.

*Immunolabelling* procedures were adapted from a previously published protocol (33) using primary antibodies directed against Tyrosine Hydroxylase (TH) (Millipore MAB318, 1:400) and ΔFosB (Abcam AB11959, 1:500). TH immunolabelling was completed using a fluorophore-coupled secondary antibody (Alexa Fluor 647, Invitrogen A21235, 1:1000), while ΔFosB required DAB revelation for best results (BA-2000 secondary antibody, 1:250, and PK6100 kit from Vector Laboratories) (**Fig. S2A,C**).

*In situ hybridization (ISH)* was performed with antisense digoxygenin-labeled complementary RNA probes designed to recognize *Gpr88*, 67-kDa glutamate decarboxylase (*Gad67*), proenkephaline (*Penk*), prodynorphine (*Pdyn*) mRNAs as previously described (23) (**Fig. S2B, D**). A new probe was also designed following the same procedures to target regulator of G-protein signaling 4 (*Rgs4*) mRNA (GenBank accession number: NM017214, targeted nucleotides: 325 to 604).

#### Digitization and semi-quantitative analysis

Slides were then digitized using the Axio Scan.Z1 and ZEN software (Zeiss, Oberkochen, Germany). The resulting images were exported for processing in Fiji (NIH, Bethesda, MD, USA) (34). As fluorescent and colorimetric stainings are not stoichiometrically related to biological content, the signal intensity was not quantified. Instead, a threshold was determined using control slides (secondary antibody alone/sense probe) or control areas within a slice (corpus callosum), and applied to all of the images from a same experiment. To evaluate the loss of dopaminergic terminals, the TH-positive signal was then quantified in each striatal area (DLS, DMS, ventral striatum) of every lesioned rat according to previously published methods(35). Regarding the nuclear markers (ΔFosB and all of the ISH targets), a fixed-size region of interest was drawn in the DLS and DMS, and the total signal-positive area was quantified (**see Fig.S2E**). For each striatum, the signal was measured over at least 3 anteroposterior locations between AP +0,2 mm and +1,2 mm, and averaged. The values were then normalized to those obtained in control rats (SHAM + miR-neg). A Fiji macro script was written to automatize the process. As the lesion and transduction extent varied, each striatum (2/brain) was considered as a biological replicate for statistical analyses.

### Statistical Analyses

Data from the experiments were analyzed using the Prism 6.0 software (GraphPad Software Inc, La Jolla, CA, USA). Different tests were performed depending on the nature of the data and the driving hypotheses that were exposed in the “study design” section. For instance, regarding behavioral data, to assess the effects of the lesion (hypothesis 1), the two groups (SHAM-neg *vs* 6OHDA-neg) were compared using two-tailed unpaired t-tests, followed by Welch corrections. Then, to evaluate the effects of *Gpr88-*KD in 6OHDA-lesioned animals (hypothesis 2), the 6OHDA + miR-*Gpr88* groups (DLS and DMS) were compared to the 6OHDA + miR-neg animals using one-way ANOVAs followed by Dunnett multiple comparison tests, as the data distribution passed normality tests. However, some of the actimeter, PPI and SND data required the use of 2-way ANOVAs followed by Sidak or Dunnett corrections where appropriate to assess interactions with independent variables (time/prepulse intensity/novelty status). Finally, as each striatal area was differentially affected by the 6OHDA lesion, the data from the immunolabelling and ISH experiments were analyzed using multiple t-tests followed by Holm-Sidak corrections.

## RESULTS

### 6OHDA stereotaxic injections induce a dopaminergic denervation limited to the DLS

In order to reproduce the loss of dopaminergic projections to sensorimotor territories of the striatum (posterior putamen) observed in early PD, we stereotaxically injected the retrograde toxin 6OHDA in the corresponding area in adult male rats (DLS) (36). Control animals (SHAM) were injected with a vehicle solution. The toxin induced a partial loss of dopaminergic afferences to the DLS, as indicated by a 48% mean reduction of Tyrosine Hydroxylase immunoreactivity over at least 1mm on the anterior-posterior axis [p<0.0001, multiple t-tests followed by Holm-Sidak corrections; **Fig. 1A, B**]. The neighboring associative (DMS) and limbic (VS) striatal areas were however not significantly affected by the lesion [in the DMS p=0.088; in the VS p=0.671; **Fig. 1B**].

**Figure 1.**
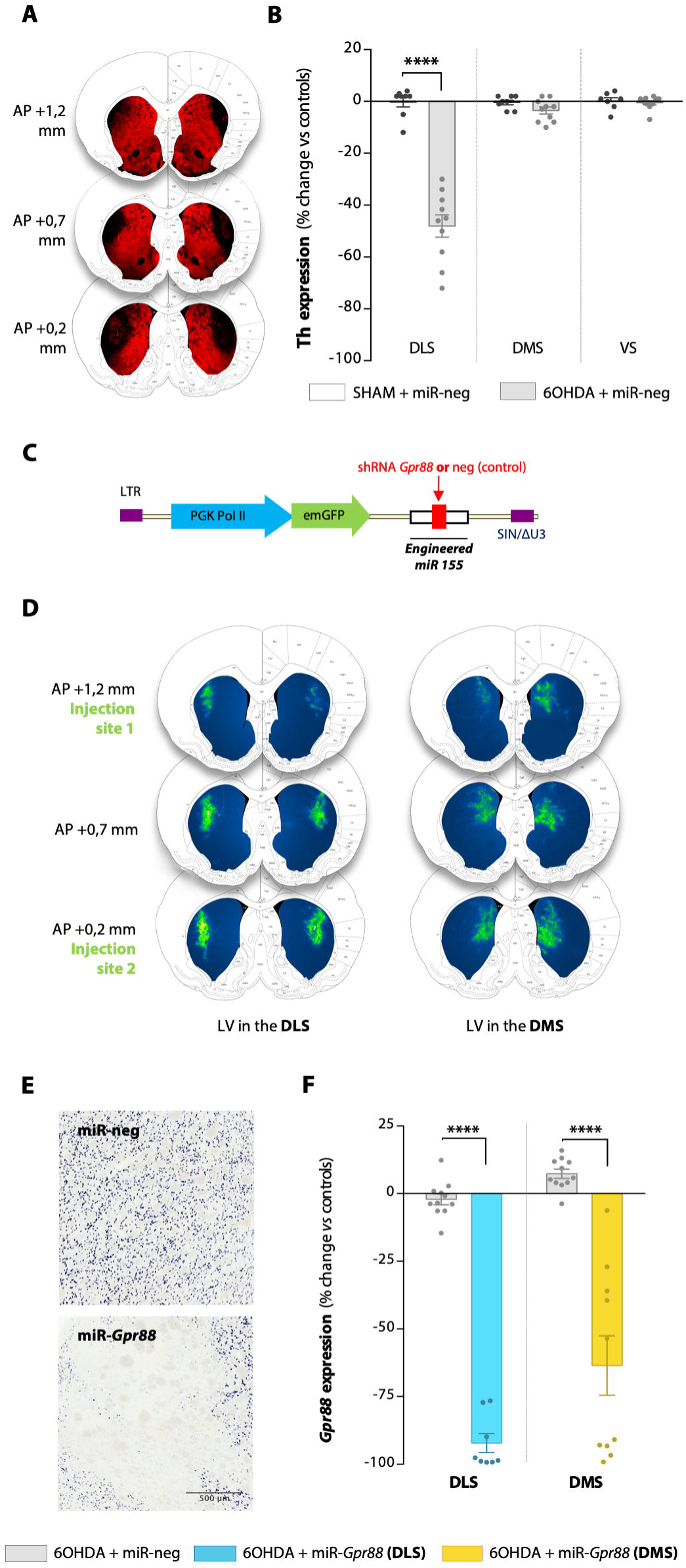
Stereotaxic injections induce a restricted loss of nigrostriatal dopaminergic projections and an efficient knock-down of Gpr88 expression in discrete striatal areas. **(A**) Representative images of 6-hydroxydopamine (6OHDA)-lesioned striata. Coronal sections stained for Tyrosine hydroxylase (TH) by immunofluorescence (+1,2 to +0,2 mm anterior to bregma), superimposed on the Paxinos and Watson rat brain atlas (29) for anatomical reference. **(B)** Quantification of the extent of Th signal loss in the dorsolateral (DLS), dorsomedial (DMS), and ventral striatum (VS). Compared to SHAM-injected rats, the 6OHDA induced a mean loss of 48% of the TH signal in the DLS, without affecting the DMS and VS. **(C)** Schematic representation of the lentiviral construct used to transduce the targeted striatal regions, two weeks after the 6OHDA/SHAM injection. **(D)** Representative images of DLS or DMS transduced striata. The lentiviruses were injected at two coordinates anterior to bregma: +1,2 and +0,2mm. For visibility purposes, a LUT was applied to the images in ImageJ in order to visualize the GFP fluorescence on a blue background. The photographs were then superimposed on the Paxinos and Watson rat brain atlas (29) for anatomical reference. **(E)** *In situ* hybridization shows a strong suppression of *Gpr88* expression induced by miR-*Gpr88*, but not miR-neg. **(F)** Quantification of the *in situ* hybridization signal for *Gpr88* mRNA following miR-neg or miR-*Gpr88* injections in the DLS or DMS. The values were normalized to those of the control group (SHAM + miR-neg). They are presented as mean ± SEM and were compared with Holm-Sidak corrected multiple t-tests. **** p<0.0001.

### Lentiviral-mediated knock-down efficiently reduces *Gpr88* expression

Two weeks after the first surgical procedure, lentiviral vectors containing a control miR (miR-neg) or a miR directed against *Gpr88* (miR-*Gpr88*) were then stereotaxically injected at multiple sites either in the DLS or the DMS of 6OHDA-lesioned rats (**Fig. 1C, D**). SHAM-lesioned rats were injected with miR-neg containing vectors. Expression of *Gpr88* mRNA was efficiently reduced in the transduced areas [p<0.0001 in DLS and DMS, multiple t-tests followed by Holm-Sidak corrections; **Fig. 1E, F**].

### Partial dopaminergic depletion of the DLS reproduces psychiatric symptoms of PD without inducing motor deficits

In order to assess whether the limited loss of DA may have an impact on locomotion and thus introduce a confounding factor when evaluating other behavioral parameters, motor behavior was assessed at two and four weeks after 6OHDA injections by measuring horizontal movements in the Actimeter. This parameter was unaffected by the lesion at both timepoints **(Fig. 2A, Fig. S3A**), as expected with a restricted lesion (35). However, although stereotyped behavior in the Actimeter test was also preserved at both timepoints, the number of rearings was reduced at 4 weeks after the lesion [2W-ANOVA: interaction of treatment x time, F(2, 44) = 4.66, p=0.015; **Fig. 2A**], indicating a decrease in novelty exploration behavior (37).

**Figure 2.**
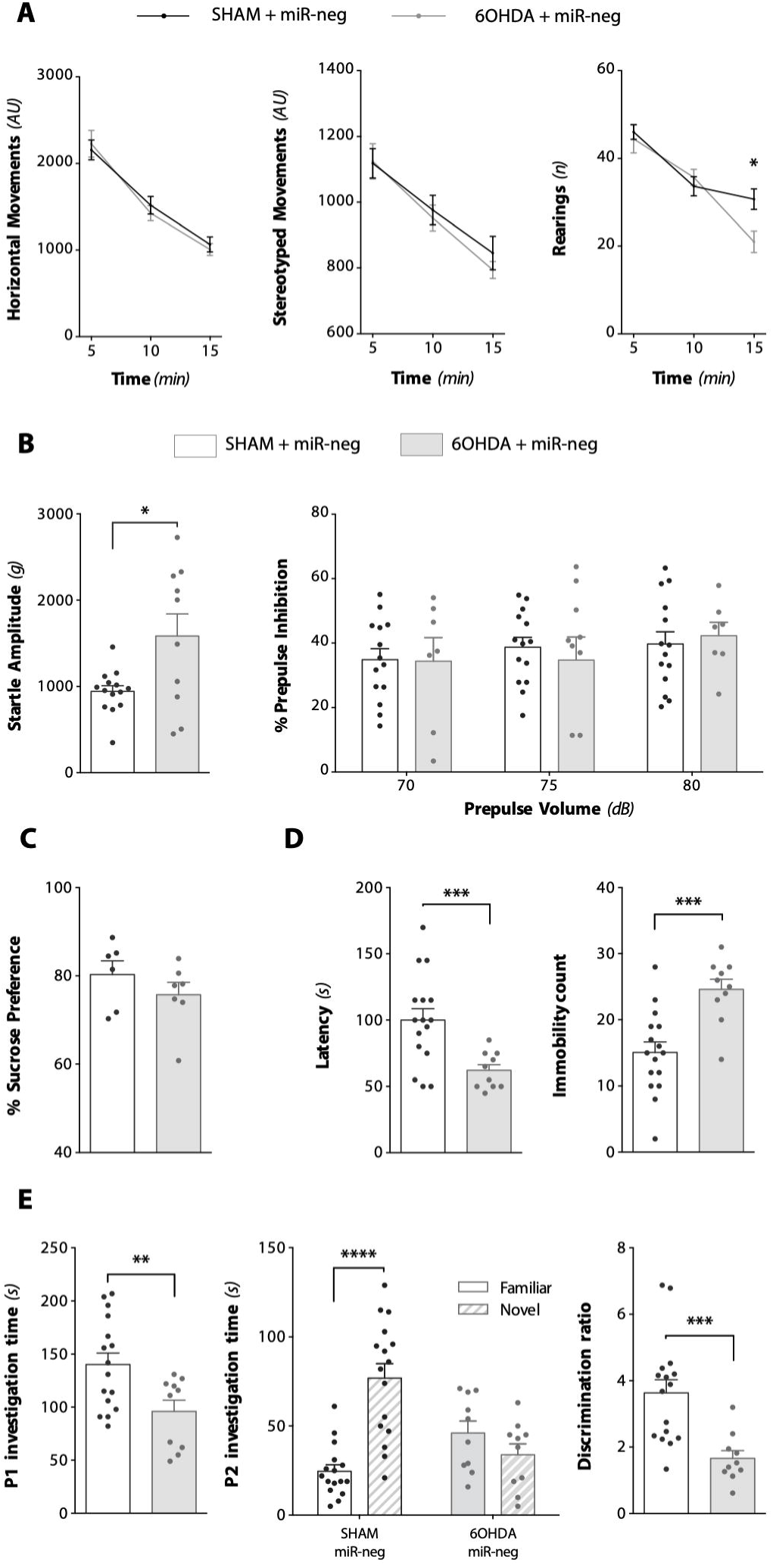
Dopamine loss in the dorsolateral striatum (DLS) reproduces psychiatric symptoms of PD. **(A)** Horizontal, stereotyped and rearing behaviors were measured with the Actimeter test 4 weeks after the 6-hydroxydopamine (6OHDA) lesion. The loss of dopamine (DA) only affected rearing at the 15-minute timepoint. **(B)** The lesion also increased the startle amplitude in response to a loud stimulus (110dB), without however affecting % Prepulse Inhibition at any of the tested prepulse intensities. **(C)** Sucrose preference was not affected by the 6OHDA lesion. **(D)** The lesion nevertheless induced a depressive-like behavior, reflected by a decrease in latency and an increase in the immobility count. **(E)** The 6OHDA lesion also reduced social interaction duration, abolished preferential interaction with a novel juvenile, and impaired the novelty discrimination ratio. Data are presented as mean ± SEM. When two groups were compared, two-tailed, Welch-corrected t-tests were performed. In the case of interactions with additional factors, two-way ANOVAs, followed by Sidak’s multiple comparison test were used (Actimeter data, %PPI, and P2 investigation time). *p<0.05, **p<0.01, ***p<0.001, ****p<0.0001.

We then evaluated sensorimotor gating, a parameter that is altered in many basal ganglia disorders including PD, and which has been linked to cognitive dysfunction in patients (38). To do so, we used the Prepulse Inhibition (PPI) test, which has previously been shown to be sensitive to 6OHDA lesions (39). Although the partial lesion of the DLS increased the startle reaction in response to a loud auditory stimulus (110dB) [P=0.037, Welch-corrected t-test; **Fig. 2B**], the level of sensorimotor gating (%PPI) was preserved at all levels of prepulse intensity used, suggesting the presence of efficient compensatory mechanisms.

Affective parameters relating to anhedonia and depression, both frequently reported in PD (3), were then assessed with the sucrose preference and forced swim tests (FST) respectively. Sucrose preference as well as general consummatory behavior, measured by daily food and water intake, were not affected by the 6OHDA lesion (**Fig. 2C, Fig. S3B**). The lesion however decreased the latency to immobility and increased immobility time in the FST [p<0.001 in both cases, Welch-corrected t-tests; **Fig. 2D**], representative of a despair-like behavior associated with depression (32).

Finally, to evaluate deficits in motivation and selective attention (a core feature of PD executive dysfunction) (40), the social interaction and novelty discrimination test was used. Loss of dopamine in the DLS decreased interaction time in the first part of the test [p=0.007, Welch-corrected t-test; **Fig. 2E**], an ecological indicator of apathy (41). Then, in the discrimination task, the lesioned animals also failed to preferentially direct their attention to the novel juvenile, which resulted in a decreased discrimination ratio [p<0.001, Welch-corrected t-test; **Fig. 2E**]. This effect was however not due to a decrease in total social interaction during the discrimination task (**Fig. S3C**), thus excluding a direct interference from the decreased motivational state.

Taken together, these behavioral results indicate that the loss of DA restricted to the DLS was sufficient to reproduce alterations relating to depression, apathy and attentional deficits, which are three of the most prevalent symptoms at the time of PD diagnosis (42).

### *Gpr88* knock-down in the DMS, but not the DLS, reverses the behavioral deficits

Although *Gpr88-*KD had no effect on horizontal and stereotyped activity in the Actimeter test, inactivation in the DMS reversed the deficit in rearing behavior at the 15-minute timepoint [p=0.020, Dunnett multiple comparison test; **Fig. 3A**]. It also increased sensorimotor gating without affecting startle amplitude [2W-ANOVA: effect of treatment on % PPI, F(2,78) = 9,79, p<0.001; **Fig. 3B**].

**Figure 3.**
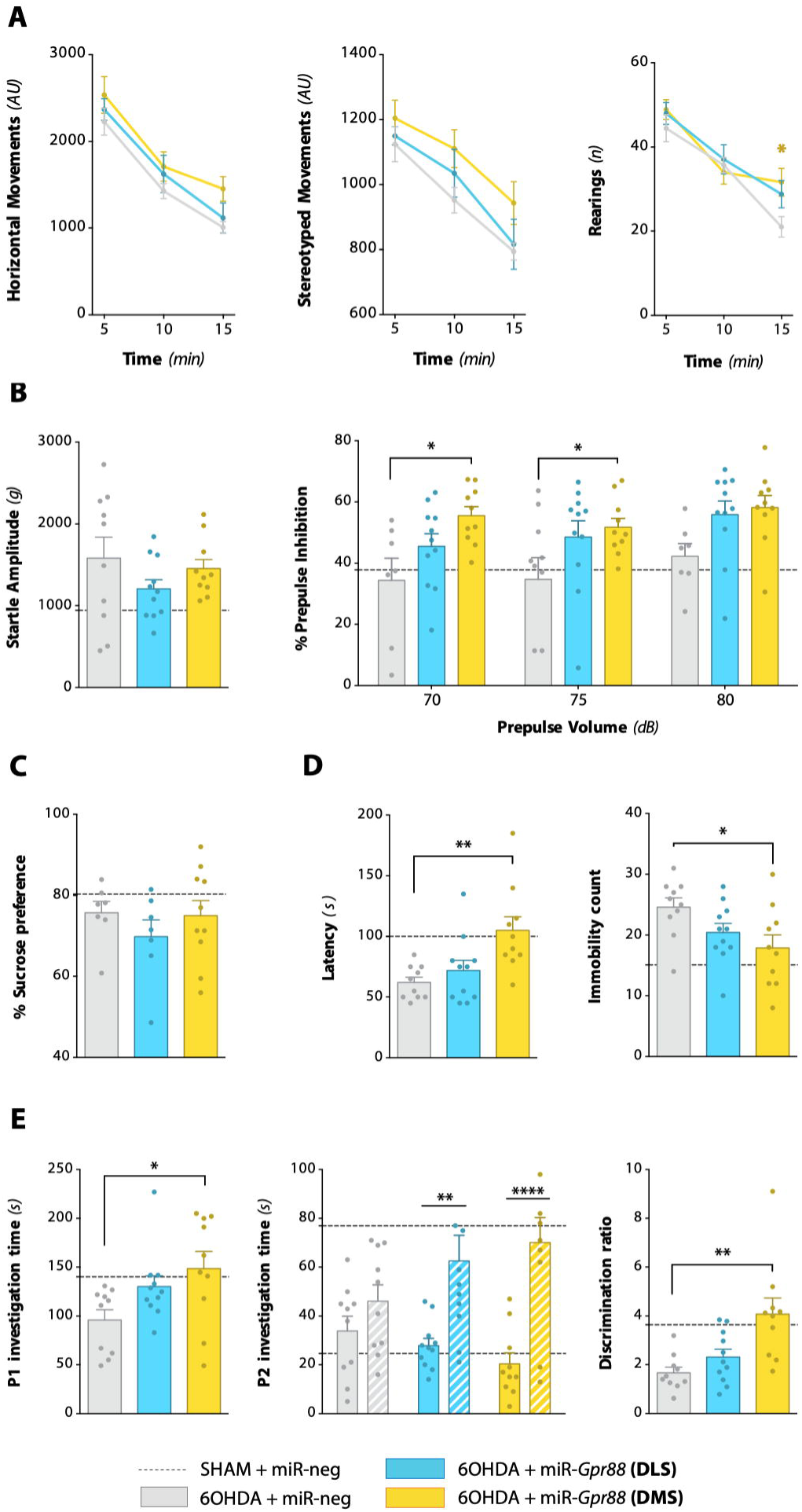
Gpr88 knock-down in the dorsomedial (DMS), but not the dorsolateral striatum (DLS), reverses the behavioral deficits. **(A)** *Gpr88-*KD in the DMS reversed the rearing deficit at the 15-minute timepoint, without affecting horizontal or stereotyped behavior. **(B)** While *Gpr88-*KD did not affect startle amplitude, it increased the % prepulse inhibition at the lower prepulse volumes**. (C)** Sucrose preference was not affected by *Gpr88-*KD. **(D)** *Gpr88-*KD in the DMS had an antidepressant-like effect as it increased latency and decreased the immobility count in the FST**. (E)** *Gpr88-*KD in the DMS increased social interaction duration, and restored novelty preference and discrimination ratio. *Gpr88-*KD in the DLS selectively increased novelty preference, but the effect was not strong enough to restore the discrimination ratio. Data are presented as mean ± SEM. For reference, a dashed horizontal line indicates the values from the control group (SHAM + miR-neg), that were presented in Fig. 2. When the three 6-hydroxydopamine (6OHDA) groups were compared, one-way ANOVAs followed by Dunnett’s multiple comparisons test were performed. In the case of interactions with additional factors, two-way ANOVAs, followed by the Dunnett multiple comparisons test were used (Actimeter data, %PPI, and P2 investigation time). *p<0.05, **p<0.01, ***p<0.001, ****p<0.0001.

Furthermore, whereas *Gpr88-*KD in either area had no impact on sucrose preference, inactivation in the DMS had an antidepressant-like effect in the FST, as it increased latency and reduced immobility time [p=0.003 and p=0.021 respectively, Dunnett multiple comparison test; **Fig. 3D**]. This reduction was mediated by an increase in swimming behavior specifically [p=0.028, Dunnett multiple comparison test; **Fig. S4A**]. No effect on consummatory behavior was observed (**Fig. S4B**).

*Gpr88-*KD in the DMS also had a pro-motivational effect as it increased social interaction duration [p=0.019, Dunnett multiple comparison test; **Fig. 3E**]. Finally, whereas the inactivation in both the DLS and DMS increased preferential interaction with the novel juvenile in the social discrimination task [p=0.004 and p<0.0001, respectively, Sidak multiple comparison test; **Fig. 3E**], only the effect in the DMS was large enough to restore the discrimination ratio [p=0.001, Dunnett multiple comparison test; **Fig. 3E**]. This pro-attentional effect was however not associated with an increase in total interaction time during the discrimination task (**Fig. S4C**).

To summarize, *Gpr88* inactivation in the intact associative DMS, but not the lesioned sensorimotor DLS, was thus able to significantly reduce the behavioral deficits that characterized this model of the psychiatric symptoms of PD.

### DA loss in the DLS induces molecular alterations in the adjacent DMS

We then assessed the expression of molecular markers of neuronal activity that are known to be modulated by 6OHDA lesions. For instance, *Prodynorphin* (*Pdyn*) and *Proenkephalin* (*Penk*) are expressed in direct and indirect pathway MSN (dMSN, iMSN) respectively, where their expression level reflects neuronal activity (43). We also evaluated the expression of *Glutamate decarboxylase 67* (*Gad67*), an enzyme involved in the synthesis of GABA, as it is considered a proxy for the global level of GABAergic transmission in the striatum, arising from MSN and interneurons (44). Strikingly, we found that despite the loss of DA being strictly limited to the DLS, significant alterations in the expression of these markers were also observed in the adjacent, intact DMS. Indeed, while the lesion expectedly decreased *Pdyn* and increased *Penk* expression levels in the DLS [p<0.0001 in both cases; multiple t-tests followed by Holm-Sidak corrections, here and throughout the next sections; **Fig. 4A**] (43), the expression of both markers was upregulated in the un-lesioned DMS [p=0.020 and p=0.013 respectively; **Fig. 4A**], indicating a local hyperactivity of dMSN and iMSN. Furthermore, *Gad67* expression was also significantly upregulated throughout the dorsal striatum [p<0.001 in the DLS, p=0.007 in the DMS; **Fig. 4A**].

**Figure 4.**
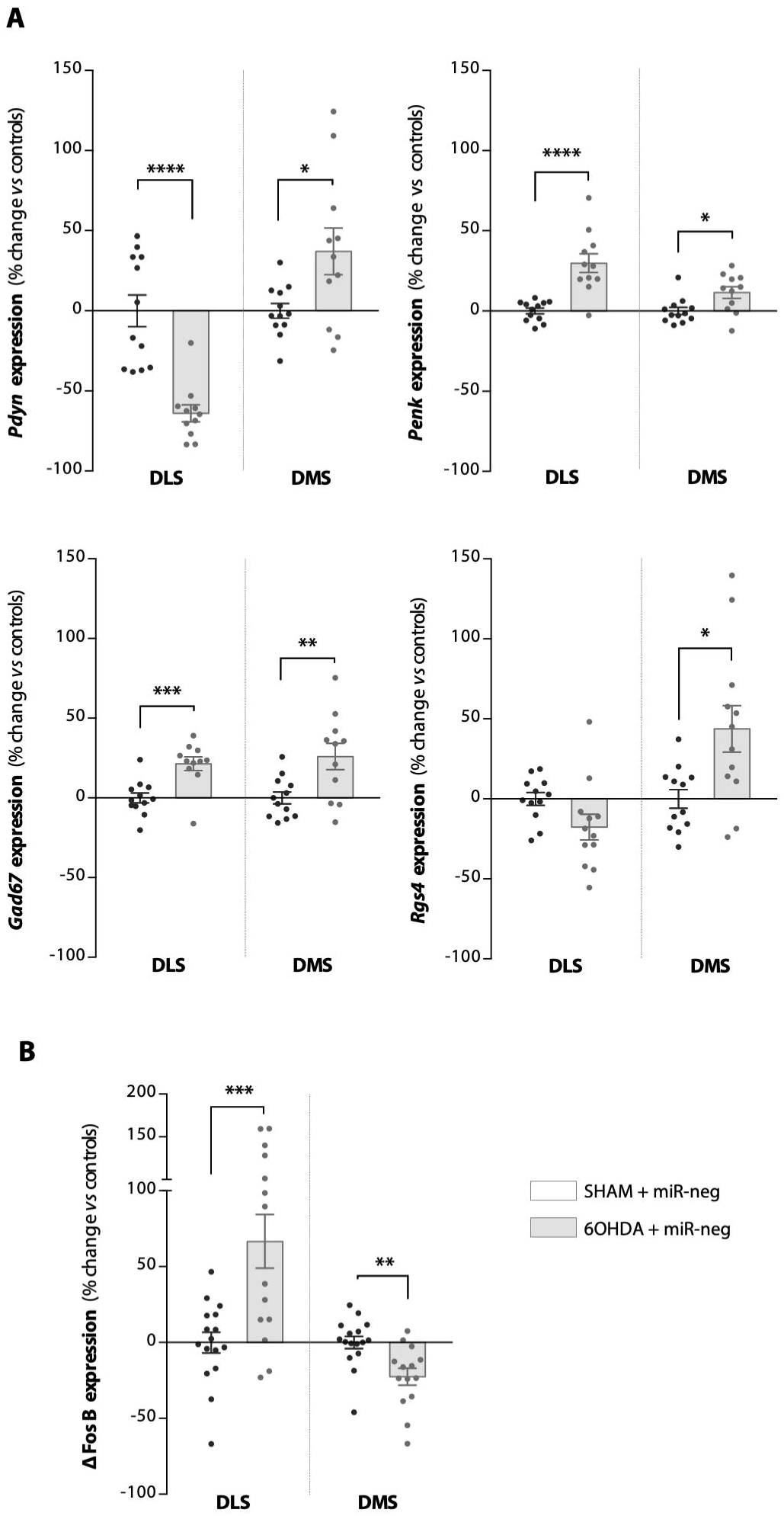
Molecular changes induced by dopamine loss in the DLS also affect the adjacent DMS. **(A)** The lesioned rats had an increased expression of *Gad67* and *Penk* in both the dorsolateral (DLS) and dorsomedial striatum (DMS). However, whereas *Pdyn* expression was strongly decreased in the lesioned DLS, it was increased in the intact DMS. The loss of dopamine (DA) also resulted in an increased *Rgs4* expression in the DMS. **(B)** The 6-hydroxydopamine (6OHDA) lesion induced a local increase in ΔFosB expression, but a decrease in the un-lesioned DMS. The values were normalized to those of the control group (SHAM + miR-neg). Data are presented as mean ± SEM, and were compared using multiple t-tests with Holm-Sidak corrections. *p<0.05, **p<0.01, ***p<0.001, ****p<0.0001.

Then, to investigate the effects of the lesion on intracellular signaling, we chose to focus on the regulator of G-protein signaling 4 (RGS4), which is involved in the pathophysiology of PD (45–47) and causally linked to GPR88 function (18). While the 6OHDA lesion induced a trend towards a decrease in the expression of *Rgs4* in the DLS, it significantly increased it in the DMS [p=0.062 and p=0.011, respectively; **Fig. 4A**]. *Gpr88* expression remained however unaffected by the loss of DA in either striatal area (**Fig. S5**).

ΔFosB, a transcription factor involved in neuropsychiatric disorders including PD (48), was then chosen as a final molecular readout, as its unique accumulation profile confers it a long-lasting effect on the regulation of striatal gene networks (49). We found that while ΔFosB expression was locally increased by the loss of DA, it was also significantly decreased in the un-lesioned DMS [p<0.001 and p=0.002, respectively; **Fig. 4B**].

Taken together, these results thus indicate that the restricted loss of Substantia Nigra DA inputs to the sensorimotor territories of the striatum alters the expression of molecular markers of neuronal activity, signaling and transcription beyond the lesioned area, significantly affecting associative territories.

### *Gpr88* inactivation in the DMS reverses alterations in *Rgs4* and **Δ**FosB expression

We then analyzed the effect of *Gpr88-*KD on the aforementioned molecular markers, and found that *Gpr88-*KD strongly reduced the expression of *Gad67* and *Rgs4* in the DLS and DMS [in the DLS: p<0.0001 for both mRNAs, in the DMS: p<0.001 for *Gad67* and p=0.002 for *Rgs4*; **Fig. 5A**]. In the DLS, *Gpr88-*KD also significantly decreased the expression of *Penk*, suggesting a reduction of iMSN hyperactivity, and induced a trend towards an increase in *Pdyn* expression [p<0.0001 and p=0.058, respectively, **Fig. 5A**]. Interestingly, *Gpr88-*KD had opposing effect on ΔFosB expression in the DLS and DMS, reverting in both cases the 6OHDA-induced alterations [p<0.001 and p=0.002, respectively, **Fig. 5B**].

**Figure 5.**
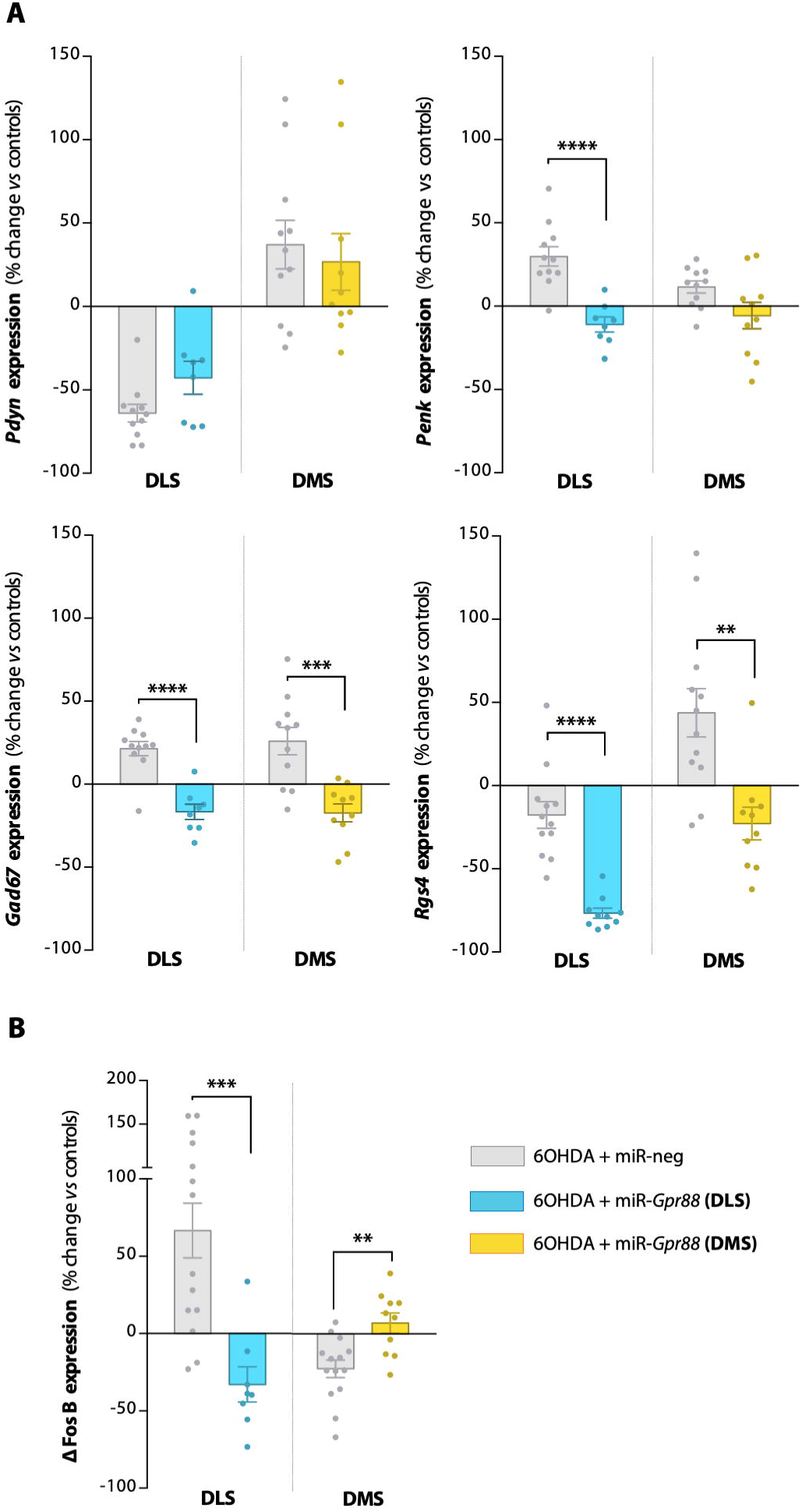
Gpr88 knock-down modulates markers of neuronal activity, signaling and transcription. **(A)** *Gpr88-*KD decreased *Gad67*, *Penk* and *Rgs4* expression levels. It however did not significantly affect *Pdyn* expression. **(B)** *Gpr88-*KD reversed ΔFosB expression in a dopamine-dependent manner (dorsolateral/dorsomedial striatum). The values were normalized to those of the control group (SHAM + miR-neg). They are presented as mean ± SEM, and were compared using multiple t-tests followed by Holm-Sidak corrections. **p<0.01, ***p<0.001, ****p<0.0001.

## DISCUSSION

In our experiments, we showed that reproducing early PD DA loss in the sensorimotor striatum (DLS) of rats induced behavioral deficits that are representative of some of the most frequent psychiatric symptoms of the disease, without however altering motor behavior. For instance, the DA loss increased immobility time in the forced swim test, an indicator of behavioral despair linked to depression (32). It also induced a decrease in social interaction and rearing behavior (related to novelty exploration (37)), two parameters that have been proposed as ecological and translational measures of apathy (41). These results are also coherent with the observations that behavioral and social aspects of apathy are the most affected in PD, and that they are specifically associated with depression (50, 51). Furthermore, PD apathy is frequently associated with cognitive impairment and dementia (4). In this regard, we also observed that the lesioned animals performed poorly in the social novelty discrimination task, suggesting an impairment of selective attention (30), a core component of the dysexecutive syndrome experienced by patients (12).

We however did not replicate deficits relating to anhedonia and sensorimotor gating that had previously been reported in other 6OHDA models of the disease using the sucrose preference and prepulse inhibition (PPI) tests (39, 52). Anhedonia is frequently reported in PD, but has however been associated with the mesolimbic dopamine pathway (53), which we did not affect in our study. Previous studies have also found that extensive dopamine loss is required to induce PPI alterations (39). In our experiments, while the lesioned rats exhibited a significantly increased startle reaction, they maintained a normal level of %PPI, suggesting the presence of efficient compensatory mechanisms. It is thus likely that the lack of PPI and sucrose preference alterations in our model is due to the restricted nature of the lesion.

Importantly, we showed that *Gpr88* inactivation in the associative (DMS), but not the sensorimotor (DLS) striatum, had pro-motivational, pro-attentional and antidepressant effects, as it reversed the behavioral deficits induced by the DA loss, without however affecting motor behavior. *Gpr88-*KD in the DMS also increased sensorimotor gating in the PPI test, indicating an increased efficiency of pre-attentive information processing, which could underlie the pro-attentional effect in the social novelty discrimination task. Interestingly, computational models of PD psychosis have linked the emergence of hallucinations to deficits in such gating and attentional processes (6), suggesting that *Gpr88-*KD may constitute a relevant target to evaluate in animal models of this symptom.

Our molecular investigations then allowed us to identify markers that were involved in both the effect of DA loss and of *Gpr88-*KD. First of all, while a previous study had reported that 6OHDA lesions induced a decrease in *Gpr88* mRNA and protein expression after 1 and 4 weeks (16), we did not observe such an effect at 6 weeks, suggesting a time-dependent recovery of *Gpr88* expression following DA loss.

At the level of neuronal activity markers, we observed that the lesion locally induced a decrease in *Pdyn* and an increase in *Penk* and *Gad67* expression levels, reflecting the imbalance in neuronal activity that has previously been characterized in PD models (43, 44). However, this restricted loss of DA also led to molecular changes in the neighboring associative territory of the striatum (DMS). Indeed, in the DMS, all of the aforementioned markers were significantly elevated, suggesting a compensatory overactivity of neurons in response to DA depletion of the DLS.

We found that the effects of *Gpr88-*KD in the lesioned DLS were mediated by a significant decrease in the expression of *Penk* and *Gad67*. This result indicates that *Gpr88-*KD preferentially affects iMSN, which is coherent with the receptor’s relative enrichment in this neuronal type (16), and the hypersensitivity to D2 agonists that has been consistently reported in *Gpr88*-KO mice (18, 20). As GPR88 is also known to modulate enkephalin receptor DOR signaling (19), the reduction in *Penk* expression we observed following *Gpr88-*KD may thus partly result from a potentiation of this receptor’s effects. In the DMS, *Gpr88-*KD reduced *Gad67* overexpression without however significantly affecting *Penk* or *Pdyn,* suggesting a potential decrease in GABA interneuron activity (44), in which GPR88 expression has been reported (16).

We then investigated *Rgs4* as a marker of intracellular signaling. RGS4 is a GTPase accelerating protein which negatively regulates the signaling of G_i/o_- and G_q_-associated receptors that have been involved in PD pathophysiology such as D2/D3 (54), mGluR 5(55) and M4 (46). Previous studies have reported slight decreases in striatal *Rgs4* expression in patients (45) and in rodents following 6OHDA injections (56). In our model, while we did observe a trend towards a decreased expression of *Rgs4* in the lesioned DLS (p=0.06), it was also strikingly elevated in the DMS. This overexpression was however completely reversed by *Gpr88-*KD, an effect that is in line with findings from KO mice (18, 19). Interestingly, in models of PD, inhibition or loss of *Rgs4* has been shown to have potent anti-parkinsonian effects (57), notably by restoring LTD in iMSN (47). Although these reported effects have been encouraging, translation to the clinic has been impeded by the high risk of side-effects due to RGS4’s ubiquitous presence (58). Targeting GPR88 may thus constitute a unique strategy to bypass these limitations by inhibiting RGS4 with high spatial relevance, as GPR88 is exclusively expressed in the CNS, and is extremely enriched in the striatum, where its expression levels match the typical pattern of dopamine loss (59).

Finally, we observed contrasting effects of the lesion and *Gpr88* inactivation on the expression of transcription factor ΔFosB. For instance, while ΔFosB accumulation in PD has previously been linked to the effects of L-DOPA treatment (48, 60), we also observed an increase in the DLS of the lesioned rats. This unexpected effect could be due to stimulation of hypersensitive D1 receptors in the DLS, resulting from a “volume transmission” effect from the un-lesioned striatum (61). In the DMS however, there was a significant decrease in ΔFosB expression, which was reversed by *Gpr88-*KD. Previous studies have shown that in the dorsal striatum, D2 stimulation increases ΔFosB accumulation, which in turn sensitizes the D1 receptor (62–64). One could thus speculate that the increased expression of ΔFosB we observed in the DMS following *Gpr88-*KD may result from a potentiation of D2 signaling through *Rgs4* inhibition. Furthermore, ΔFosB-mediated D1 sensitization may thus potentiate dMSN signaling, and help promote goal-directed behavior. For example, the antidepressant effect of *Gpr88-*KD observed in the FST may be due to its modulation of ΔFosB, as fluoxetine is also known to increase the expression of this transcription factor in the DMS (65). Nevertheless, the fact that *Gpr88-*KD had an opposing effect on ΔFosB expression in the DLS suggests a complex interaction with dopamine signaling, which deserves to be further investigated.

To conclude, our results support the relevance of GPR88 as a therapeutic target for the psychiatric symptoms of PD, as we demonstrate in our model that knocking-down its expression in the associative striatum reduces the behavioral deficits and normalizes the expression of *Rgs4* and of ΔFosB, which have both been previously linked to PD pathophysiology.

Beyond supporting GPR88 as a potential therapeutic target, this study may also provide insights regarding the etiology of the psychiatric symptoms of PD. Indeed, while these symptoms have been linked to alterations in multiple neurotransmitter systems (26), clustering studies of *de novo* patients suggest that the loss of striatal dopamine in particular may play an important role in their emergence (66, 67). This idea is further supported by studies in rodent models, which have consistently shown that inducing extensive DA loss throughout the striatum could induce representative behavioral deficits (52). Nevertheless, these models haven’t necessarily accounted for the frequent pre-motor appearance of these symptoms, when the DA loss is mainly limited to sensorimotor areas of the striatum (25). In this regard, our results show that such a restricted loss of dopamine is *per se* sufficient to reproduce these behavioral alterations in an animal model.

These psychiatric aspects of PD have however typically been associated with DA dysfunctions in associative networks (DMS) (26), which are relatively preserved in early stages of the disease as well as in our model. Nevertheless, we did observe an overexpression of neuronal activity markers in the DMS, suggesting possible interactions between striatal areas. Interestingly, fMRI studies of PD patients have also found that while functional connectivity decreased in the dopamine-deprived sensorimotor networks of the basal ganglia, increases could also be observed in associative networks (68–70). These increases are thought to reflect early PD allostatic mechanisms, by which associative networks may expand or strengthen their connectivity to compensate for the loss of DA in sensorimotor territories. Although the underlying mechanisms are currently unknown, recently characterized striato-nigro-striatal cellular pathways may provide a system for such lateral transfer of information (71). However, due to the resulting overlap of networks within limited striatal computational resources, it has been suggested that this compensatory mechanism could lead to a “neural bottleneck” or “overload” of associative networks, and impairment in related behaviors (36, 68, 69). This hypothesis could thus explain the emergence of psychiatric deficits relating to cognition and motivation during the earliest stages of PD and in our model of the disease, before the loss of DA affects associative networks.

In this context, the results from our investigations may suggest a molecular mechanism for the “overload” hypothesis. Indeed, as Dynorphin downregulates both nigrostriatal dopamine release and the induction of immediate early genes in MSN (such as Fos family genes) (43), *Pdyn* overexpression in the DMS of 6OHDA-lesioned rats may act as a throttle on dMSN activity and on the execution of goal-directed behaviors. Furthermore, the increased expression of *Rgs4* we observed in the DMS might also contribute to this effect by reducing the inhibitory control of G_i/o_ and G_q_ receptors over iMSN activity, which could further distort action selection and execution.

Although imbalances in DMS/DLS dynamics are emerging as key players in the pathophysiology of basal ganglia disorders such as addiction (72) and OCD (73), this study is the first to our knowledge to enquire this aspect in a rodent model of PD. Several elements nevertheless need to be further investigated. For instance, behavioral tests assessing specific cognitive and motivational parameters would be welcome, to further characterize the effects of the lesion and of *Gpr88-*KD. As the model we used recapitulates aspects of early-stage PD, it will also be necessary to assess the effects of *Gpr88-*KD in models of advanced PD, presenting more extensive DA loss and alterations in other neurotransmitter systems. Furthermore, while our molecular experiments have provided some clues regarding the mechanisms involved, additional in-depth studies are required, as very little is known about the signaling pathways regulated by GPR88 in different striatal cell types (dMSN/iMSN/interneurons).

Provided that these questions can be answered, there is still an essential issue to consider in order to advance the translational process to the clinic. Specifically, while GPR88’s restricted expression makes it an ideal therapeutic target, no antagonist has so far been developed for this orphan receptor. It is thus crucial to develop lead identification efforts. Nevertheless, gene therapy approaches have showed successful applications for PD (74). A recent study notably found that a *GAD* gene therapy approach targeting the subthalamic nucleus reduced motor impairment in patients by inducing changes in basal ganglia connectivity (75) that are reminiscent of those we observed in our model. Based on our findings, *GPR88* gene therapy could thus constitute a highly relevant non-dopaminergic strategy to treat the psychiatric symptoms of PD through striatal-level modulation of allostatic basal ganglia function.

## ACKNOWLEDGMENTS

The lentiviral vectors were produced by the in-house iVECTOR facility. *In vivo* experiments were carried out in the PHENOPARC core facility, and the histological procedures were performed at the HISTOMICS platform, both also situated within the ICM. The authors thank Annick Prigent for her time and advice regarding the histological processing of the samples.

The PHENOPARC Core is supported by 2 “Investissements d’avenir” (ANR-10-IAIHU-06 and ANR-11-INBS-0011-NeurATRIS) and the “Fondation pour la Recherche Médicale”. The research leading to these results has received funding from the program “Investissements d’avenir” ANR-10-IAIHU-06. B.G. was supported by a scholarship from the French Ministry of Research, awarded by the Brain-Cognition-Behavior Doctoral School of the Sorbonne University.

Earlier versions of the data were presented in posters at the FENS Forum 2018, the ECNP Workshop for Early Career Scientists 2018, and at the NeuroFrance 2017 Forum.

## Author contributions

R.M. supervised the project. B.G. and R.M. designed the experiments. B.G., J.P., R.M., M.I. and A.D.T. performed the stereotaxic surgeries. B.G. carried out the behavioral studies. J.P. and B.G. scored the FST videos. N.F.B., J.P., A.D.T. and B.G. undertook the histological processing and experiments. P.R. provided advice for lentiviral vector design, production and use. Statistical analyses were designed and executed by B.G. The manuscript was drafted by B.G. and R.M., and was reviewed by all authors.

## REFERENCES

1. Duncan GW, Khoo TK, Yarnall AJ, O’Brien JT, Coleman SY, Brooks DJ, et al. (2014): Health-related quality of life in early Parkinson’s disease: The impact of nonmotor symptoms. Mov Disord. 29: 195–202.

2. Sierra M, Carnicella S, Strafella AP, Bichon A, Lhommée E, Castrioto A, et al. (2015): Apathy and impulse control disorders: Yin & yang of dopamine dependent behaviors. J Parkinsons Dis. 5: 625–636.

3. Schapira AHV, Chaudhuri KR, Jenner P (2017): Non-motor features of Parkinson disease. Nat Rev Neurosci. 18: 435–450.

4. Pagonabarraga J, Kulisevsky J, Strafella AP, Krack P (2015): Apathy in Parkinson’s disease: clinical features, neural substrates, diagnosis, and treatment. Lancet Neurol. 14: 518–31.

5. Anang JBM, Gagnon J-F, Bertrand J-A, Romenets SR, Latreille V, Panisset M, et al. (2014): Predictors of dementia in Parkinson disease: A prospective cohort study. Neurology. 83: 1253–1260.

6. Ffytche DH, Creese B, Politis M, Chaudhuri KR, Weintraub D, Ballard C, Aarsland D (2017): The psychosis spectrum in Parkinson disease. Nat Rev Neurol. 13: 81–95.

7. Dujardin K, Langlois C, Plomhause L, Carette AS, Delliaux M, Duhamel A, Defebvre L (2014): Apathy in untreated early-stage Parkinson disease: Relationship with other non-motor symptoms. Mov Disord. 29: 1796–1801.

8. Darweesh SKL, Verlinden VJA, Stricker BH, Hofman A, Koudstaal PJ, Ikram MA (2016): Trajectories of prediagnostic functioning in Parkinson’s disease. Brain. 1–13.

9. Aarsland D, Brønnick K, Alves G, Tysnes OB, Pedersen KF, Ehrt U, Larsen JP (2009): The spectrum of neuropsychiatric symptoms in patients with early untreated Parkinson’s disease. J Neurol Neurosurg Psychiatry. 80: 928–930.

10. Rodriguez-Oroz MC, Jahanshahi M, Krack P, Litvan I, Macias R, Bezard E, Obeso JA (2009): Initial clinical manifestations of Parkinson’s disease: features and pathophysiological mechanisms. Lancet Neurol. 8: 1128–1139.

11. Pagonabarraga J, Martinez-Horta S, Fernández de Bobadilla R, Pérez J, Ribosa-Nogué R, Marín J, et al. (2016): Minor hallucinations occur in drug-naive Parkinson’s disease patients, even from the premotor phase. Mov Disord. 31: 45–52.

12. Svenningsson P, Westman E, Ballard C, Aarsland D (2012): Cognitive impairment in patients with Parkinson’s disease: Diagnosis, biomarkers, and treatment. Lancet Neurol. 11: 697–707.

13. Del Zompo M, Severino G, Ardau R, Chillotti C, Piccardi M, Dib C, et al. (2010): Genome-scan for bipolar disorder with sib-pair families in the Sardinian population: A new susceptibility locus on chromosome 1p22-p21? Am J Med Genet Part B Neuropsychiatr Genet. 153: 1200–1208.

14. Del Zompo M, Deleuze J-F, Chillotti C, Cousin E, Niehaus D, Ebstein RP, et al. (2014): Association study in three different populations between the GPR88 gene and major psychoses. Mol Genet genomic Med. 2: 152–9.

15. Alkufri F, Shaag A, Abu-Libdeh B, Elpeleg O (2016): Deleterious mutation in GPR88 is associated with chorea, speech delay, and learning disabilities. Neurol Genet. 2: e64–e64.

16. Massart R, Guilloux JP, Mignon V, Sokoloff P, Diaz J (2009): Striatal GPR88 expression is confined to the whole projection neuron population and is regulated by dopaminergic and glutamatergic afferents. Eur J Neurosci. 30: 397–414.

17. Jin C, Decker AM, Huang X-P, Gilmour BP, Blough BE, Roth BL, et al. (2014): Synthesis, Pharmacological Characterization, and Structure–Activity Relationship Studies of Small Molecular Agonists for the Orphan GPR88 Receptor. ACS Chem Neurosci. 5: 576–587.

18. Quintana A, Sanz E, Wang W, Storey GP, Güler AD, Wanat MJ, et al. (2012): Lack of GPR88 enhances medium spiny neuron activity and alters motor- and cue-dependent behaviors. Nat Neurosci. 15: 1547–55.

19. Meirsman a. C, Le Merrer J, Pellissier LP, Diaz J, Clesse D, Kieffer BL, Becker J a. J (2015): Mice lacking GPR88 show motor deficit, improved spatial learning and low anxiety reversed by delta opioid antagonist. Biol Psychiatry.. doi: 10.1016/j.biopsych.2015.05.020.

20. Logue SF, Grauer SM, Paulsen J, Graf R, Taylor N, Sung MA, et al. (2009): The orphan GPCR, GPR88, modulates function of the striatal dopamine system: A possible therapeutic target for psychiatric disorders? Mol Cell Neurosci. 42: 438–447.

21. Rainwater A, Sanz E, Palmiter RD, Quintana A (2017): Striatal GPR88 Modulates Foraging Efficiency. J Neurosci. 37: 7939–7947.

22. Maroteaux G, Arefin TM, Harsan L-A, Darcq E, Ben Hamida S, Kieffer BL (2018): Lack of anticipatory behavior in GPR88 knockout mice revealed by automatized home cage phenotyping. Genes, Brain Behav. e12473.

23. Ingallinesi M, Le Bouil L, Faucon Biguet N, Do Thi a, Mannoury la Cour C, Millan MJ, et al. (2015): Local inactivation of GPR88 in the nucleus accumbens attenuates behavioral deficits elicited by the neonatal administration of phencyclidine in rats. Mol Psychiatry. 20: 951–958.

24. Ben Hamida S, Mendonça-Netto S, Arefin TM, Nasseef MT, Boulos L-J, McNicholas M, et al. (2018): Increased Alcohol Seeking in Mice Lacking GPR88 Involves Dysfunctional Mesocorticolimbic Networks. Biol Psychiatry.. doi: 10.1016/J.BIOPSYCH.2018.01.026.

25. Brooks DJ, Pavese N (2011): Imaging biomarkers in Parkinson’s disease. Prog Neurobiol. 95: 614–628.

26. Qamar MA, Sauerbier A, Politis M, Carr H, Loehrer P, Chaudhuri KR (2017): Presynaptic dopaminergic terminal imaging & non-motor symptoms assessment of Parkinson’s disease: Evidence for dopaminergic basis? Parkinsons Dis. 3: 1–19.

27. Kilkenny C, Browne WJ, Cuthill IC, Emerson M, Altman DG (2010): Improving bioscience research reporting: The arrive guidelines for reporting animal research. PLOS Biol. 8. doi: 10.3390/ani4010035.

28. Faul F, Erdfelder E, Lang A-G, Buchner A (2007): G*Power: A flexible statistical power analysis program for the social, behavioral, and biomedical sciences. Behav Res Methods. 39: 175–191.

29. Paxinos G, Watson CRR, Emson PC (1980): AChE-stained horizontal sections of the rat brain in stereotaxic coordinates. J Neurosci Methods. 3: 129–149.

30. Terranova J-P, Chabot C, Barnouin M-C, Perrault G, Depoortere R, Griebel G, Scatton B (2005): SSR181507, a dopamine D2 receptor antagonist and 5-HT1A receptor agonist, alleviates disturbances of novelty discrimination in a social context in rats, a putative model of selective attention deficit. Psychopharmacology (Berl). 181: 134–144.

31. Valsamis B, Schmid S (2011): Habituation and prepulse inhibition of acoustic startle in rodents. J Vis Exp. 1–10.

32. Slattery D a, Cryan JF (2012): Using the rat forced swim test to assess antidepressant-like activity in rodents. Nat Protoc. 7: 1009–1014.

33. Schneider Gasser EM, Straub CJ, Panzanelli P, Weinmann O, Sassoè-Pognetto M, Fritschy J-M (2006): Immunofluorescence in brain sections: simultaneous detection of presynaptic and postsynaptic proteins in identified neurons. Nat Protoc. 1: 1887–97.

34. Schindelin J, Arganda-Carreras I, Frise E, Kaynig V, Longair M, Pietzsch T, et al. (2012): Fiji: An open source platform for biological image analysis. Nat Methods. 9: 676–682.

35. Drui G, Carnicella S, Carcenac C, Favier M, Bertrand a, Boulet S, Savasta M (2014): Loss of dopaminergic nigrostriatal neurons accounts for the motivational and affective deficits in Parkinson’s disease. Mol Psychiatry. 19: 358–367.

36. Redgrave P, Rodriguez M, Smith Y, Rodriguez-Oroz MC, Lehericy S, Bergman H, et al. (2010): Goal-directed and habitual control in the basal ganglia: implications for Parkinson’s disease. Nat Rev Neurosci. 11: 760–772.

37. Lever C, Burton S, Ο’Keefe J (2006): Rearing on Hind Legs, Environmental Novelty, and the Hippocampal Formation. Rev Neurosci. 17: 111–133.

38. Zoetmulder M, Biernat HB, Nikolic M, Korbo L, Friberg L, Jennum PJ (2014): Prepulse inhibition is associated with attention, processing speed, and 123I-FP-CIT SPECT in Parkinson’s Disease. J Parkinsons Dis. 4: 77–87.

39. Issy AC, Padovan-Neto FE, Lazzarini M, Bortolanza M, Del-Bel E (2015): Disturbance of sensorimotor filtering in the 6-OHDA rodent model of Parkinson’s disease. Life Sci. 125: 71–78.

40. Dujardin K, Leentjens AFG, Langlois C, Moonen AJH, Duits AA, Carette AS, Duhamel A (2013): The spectrum of cognitive disorders in Parkinson’s disease: A data-driven approach. Mov Disord. 28: 183–189.

41. Cathomas F, Hartmann MN, Seifritz E, Pryce CR, Kaiser S (2015): The translational study of apathy-an ecological approach. Front Behav Neurosci. 9: 241.

42. Pont-Sunyer C, Hotter A, Gaig C, Seppi K, Compta Y, Katzenschlager R, et al. (2015): The onset of nonmotor symptoms in parkinson’s disease (the ONSET PD study). Mov Disord. 30: 229–237.

43. Steiner H, Gerfen CR (1998): Role of dynorphin and enkephalin in the regulation of striatal output pathways and behavior. Exp Brain Res. 123: 60–76.

44. Consolo S, Morelli M, Rimoldi M, Giorgi S, Di Chiara G (1999): Increased striatal expression of glutamate decarboxylase 67 after priming of 6-hydroxydopamine-lesioned rats. Neuroscience. 89: 1183–1187.

45. Zhang Y, James M, Middleton FA, Davis RL (2005): Transcriptional Analysis of Multiple Brain Regions in Parkinson’s Disease Supports the Involvement of Specific Protein Processing, Energy Metabolism, and Signaling Pathways, and Suggests Novel Disease Mechanisms. Am J Med Genet Part B Neuropsychiatr Genet. 16: 5–16.

46. Ding J, Guzman JN, Tkatch T, Chen S, Goldberg JA, Ebert PJ, et al. (2006): RGS4-dependent attenuation of M4 autoreceptor function in striatal cholinergic interneurons following dopamine depletion. Nat Neurosci. 9: 832–842.

47. Lerner T, Kreitzer A (2012): RGS4 Is Required for Dopaminergic Control of Striatal LTD and Susceptibility to Parkinsonian Motor Deficits. Neuron. 73: 347–359.

48. Lindgren HS, Rylander D, Iderberg H, Andersson M, O’Sullivan SS, Williams DR, et al. (2011): Putaminal upregulation of FosB/*_Δ_*FosB-like immunoreactivity in Parkinson’s disease patients with Dyskinesia. J Parkinsons Dis. 1: 347–357.

49. Nestler EJ (2015): ΔfosB: A transcriptional regulator of stress and antidepressant responses. Eur J Pharmacol. 753: 66–72.

50. Ang Y-S, Lockwood PL, Kienast A, Plant O, Drew D, Slavkova E, et al. (2018): Differential impact of behavioral, social, and emotional apathy on Parkinson’s disease. Ann Clin Transl Neurol. 1–6.

51. Ang YS, Lockwood P, Apps MAJ, Muhammed K, Husain M (2017): Distinct subtypes of apathy revealed by the apathy motivation index. PLoS One. 12: 1–15.

52. Magnard R, Vachez Y, Carcenac C, Krack P, David O, Savasta M, et al. (2016): What can rodent models tell us about apathy and associated neuropsychiatric symptoms in Parkinson’s disease? Transl Psychiatry. 6: e753.

53. Husain M, Roiser JP (2018): Neuroscience of apathy and anhedonia: A transdiagnostic approach. Nat Rev Neurosci. 19: 470–484.

54. Min C, Cheong SY, Cheong SJ, Kim M, Cho DI, Kim KM (2012): RGS4 exerts inhibitory activities on the signaling of dopamine D 2 receptor and D 3 receptor through the N-terminal region. Pharmacol Res. 65: 213–220.

55. Schwendt M, Sigmon SA, McGinty JF (2012): RGS4 overexpression in the rat dorsal striatum modulates mGluR5- and amphetamine-mediated behavior and signaling. Psychopharmacology (Berl). 221: 621–635.

56. Geurts M, Maloteaux JM, Hermans E (2003): Altered expression of regulators of G-protein signaling (RGS) mRNAs in the striatum of rats undergoing dopamine depletion. Biochem Pharmacol. 66: 1163–1170.

57. Ko WKD, Martin-Negrier ML, Bezard E, Crossman AR, Ravenscroft P (2014): RGS4 is involved in the generation of abnormal involuntary movements in the unilateral 6-OHDA-lesioned rat model of Parkinson’s disease. Neurobiol Dis. 70: 138–148.

58. Erdely HA, Lahti RA, Lopez MB, Myers CS, Roberts RC, Tamminga CA, Vogel MW (2004): SHORT COMMUNICATION Regional expression of RGS4 mRNA in human brain y. Neuroscience. 19: 3125–3128.

59. Waes V Van, Tseng KY, Steiner H (2011): GPR88: A putative signaling molecule predominantly expressed in the striatum: Cellular localization and developmental regulation. Basal Ganglia. 1: 83–89.

60. Winkler C, Kirik D, Björklund A, Cenci MA (2002): l-DOPA-Induced Dyskinesia in the Intrastriatal 6-Hydroxydopamine Model of Parkinson’s Disease: Relation to Motor and Cellular Parameters of Nigrostriatal Function. Neurobiol Dis. 10: 165–186.

61. Taber KH, Hurley RA (2014): Volume Transmission in the Brain: Beyond the Synapse. J Neuropsychiatry Clin Neurosci. 26: 1–4.

62. Saka E, Elibol B, Erdem S, Dalkara T (1999): Compartmental changes in expression of c-Fos and FosB proteins in intact and dopamine-depleted striatum after chronic apomorphine treatment. Brain Res. 825: 104–114.

63. Crocker SJ, Morelli M, Wigle N, Nakabeppu Y, Robertson GS (1998): D1-receptor-related priming is attenuated by antisense-meditated “knock-down” of fosB expression. Mol Brain Res. 53: 69–77.

64. Wirtshafter D, Schardt G, Asin KE (1997): Compartmentally specific effects of quinpirole on the striatal Fos expression induced by stimulation of D1-dopamine receptors in intact rats. Brain Res. 771: 271–277.

65. Vialou V, Thibault M, Kaska S, Cooper S, Gajewski P, Eagle A, et al. (2015): Differential induction of FosB isoforms throughout the brain by fluoxetine and chronic stress. Neuropharmacology. 99: 28–37.

66. Erro R, Vitale C, Amboni M, Picillo M, Moccia M, Longo K, et al. (2013): The Heterogeneity of Early Parkinson’s Disease: A Cluster Analysis on Newly Diagnosed Untreated Patients. PLoS One. 8: 1–8.

67. Moccia M, Pappatà S, Picillo M, Erro R, Coda ARD, Longo K, et al. (2014): Dopamine transporter availability in motor subtypes of de novo drug-naïve Parkinson’s disease. J Neurol. 261: 2112–2118.

68. Helmich RC, Derikx LC, Bakker M, Scheeringa R, Bloem BR, Toni I (2010): Spatial remapping of cortico-striatal connectivity in parkinson’s disease. Cereb Cortex. 20: 1175–1186.

69. Sharman M, Valabregue R, Perlbarg V, Marrakchi-Kacem L, Vidailhet M, Benali H, et al. (2013): Parkinson’s disease patients show reduced cortical-subcortical sensorimotor connectivity. Mov Disord. 28: 447–454.

70. Bell PT, Gilat M, O’Callaghan C, Copland DA, Frank MJ, Lewis SJG, Shine JM (2015): Dopaminergic basis for impairments in functional connectivity across subdivisions of the striatum in Parkinson’s disease. Hum Brain Mapp. 36: 1278–1291.

71. Lerner TN, Shilyansky C, Davidson TJ, Evans KE, Beier KT, Zalocusky KA, et al. (2015): Intact-Brain Analyses Reveal Distinct Information Carried by SNc Dopamine Subcircuits. Cell. 162: 635–647.

72. Belin D, Belin-Rauscent A, Murray JE, Everitt BJ (2013): Addiction: failure of control over maladaptive incentive habits. Curr Opin Neurobiol. 564–572.

73. Burguière E, Monteiro P, Mallet L, Feng G, Graybiel AM (2015): Striatal circuits, habits, and implications for obsessive-compulsive disorder. Curr Opin Neurobiol. 30: 59–65.

74. Palfi S, Gurru J, Le H, Howard K, Ralph GS, Mason sarah, et al. (2018): Long-term follow up of a phase 1/2 study of ProSavin, a lentiviral vector gene therapy for Parkinson’s disease. Hum Gene Ther Clin Dev. 33: humc.2018.081.

75. Niethammer M, Tang CC, Vo A, Nguyen N, Spetsieris P, Dhawan V, et al. (2018): Gene therapy reduces Parkinsons disease symptoms by reorganizing functional brain connectivity. Sci Transl Med. 1–12.

