## Supplementary figures legends for "*G-protein coupled receptor 88* knock-down in the associative striatum reduces the psychiatric symptoms in a translational model of Parkinson’s disease"

*Supplementary Figure 1 – Experimental design and timeline.*

*Supplementary Figure 2 – Histological characterization.*

**(A-D)** High resolution scans of 12µm coronal slices from the same animal (6OHDA + miR-*Gpr88* in the DMS), stained for **(A)** Tyrosine Hydroxylase, **(B)** *Gpr88*, **(C)** ∆FosB, **(D)** *Gad67*, *Pdyn* and *Penk***. (E)** Summary of the image processing and quantification workflow, detailed in the methods section.

*Supplementary Figure 3 – Behavioral effects of the 6-hydroxydopamine (6OHDA) lesion: additional data.*

**(A)** The 6OHDA lesion had no effect on horizontal, stereotyped and rearing behavior at 2 weeks after injection. **(B)** The lesion did not affect consummatory behavior, **(C)** or total social investigation time during the discrimination task (P2). **(D)** Swimming, climbing and diving behaviors were also quantified during the FST. The increase in immobility count induced by the lesion (Fig. 2D) was mediated by a decrease in swimming. Data are presented as mean ± SEM. When two groups were compared, two-tailed, Welch-corrected t-tests were performed (B and C). In the case of interactions with additional factors, two-way ANOVAs, followed by Sidak’s multiple comparison test were used. *p<0.05.

*Supplementary Figure 4 – Behavioral effects of Gpr88 knock-down: additional data.*

**(A**) *Gpr88*-KD in the DMS reduced the immobility count (Fig. 3D) by specifically increasing swimming behavior. **(B)** *Gpr88*-KD did not affect consummatory behavior, **(C)** or total social interaction duration in the discrimination task. Data are presented as mean ± SEM, and were compared using one-way ANOVAs followed by Dunnett multiple comparisons tests. For reference, a dashed horizontal line indicates the values from the control group (SHAM miR-neg), that were presented in Fig. 2. *p<0.05.

*Supplementary Figure 5 – The 6OHDA lesion does not affect Gpr88 expression.*

The values were normalized to those of the control group (SHAM + miR-neg). Data are presented as mean ± SEM, and were compared using multiple t-tests with Holm-Sidak corrections.
