## Supplementary figures and images for "*G-protein coupled receptor 88* knock-down in the associative striatum reduces the psychiatric symptoms in a translational model of Parkinson’s disease"

### Supplementary figure 1

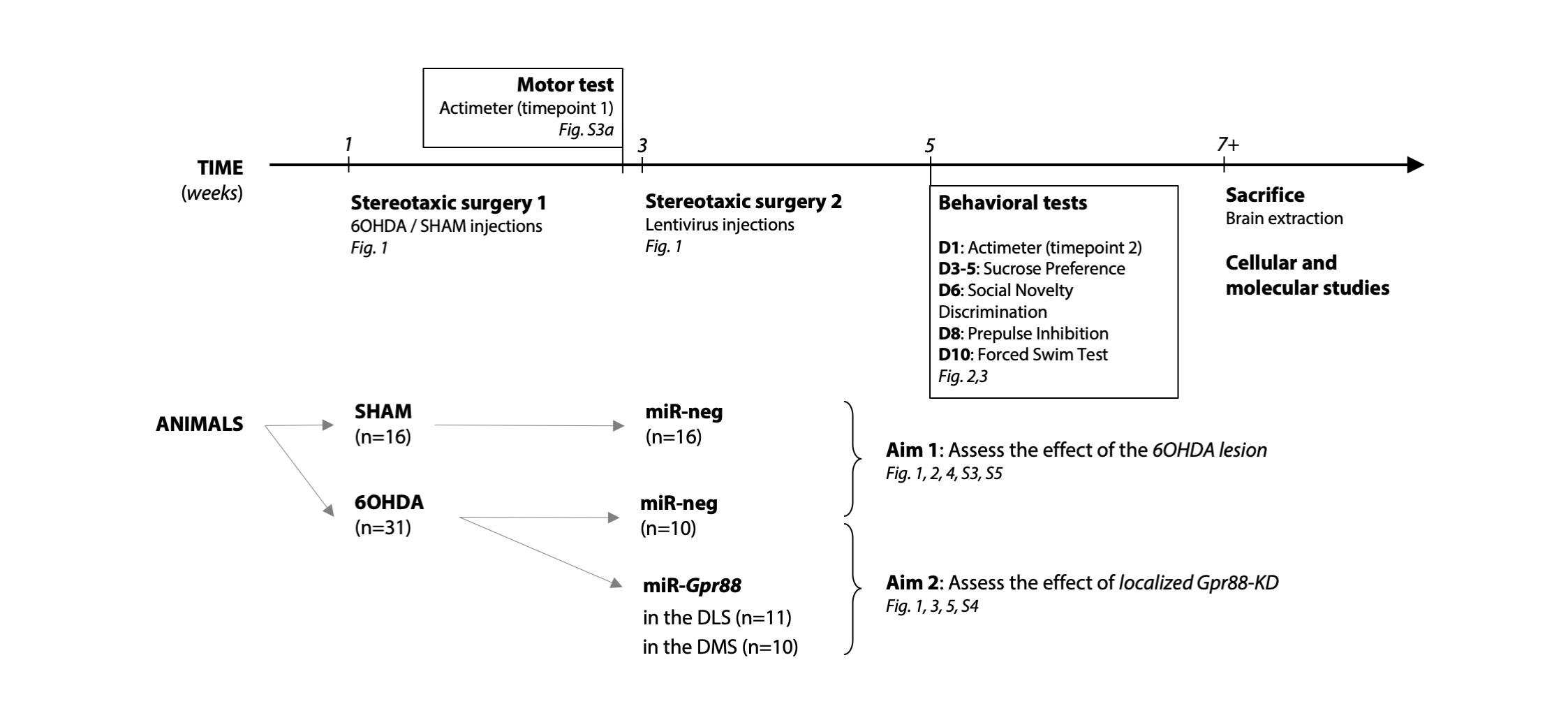

### Supplementary figure 2

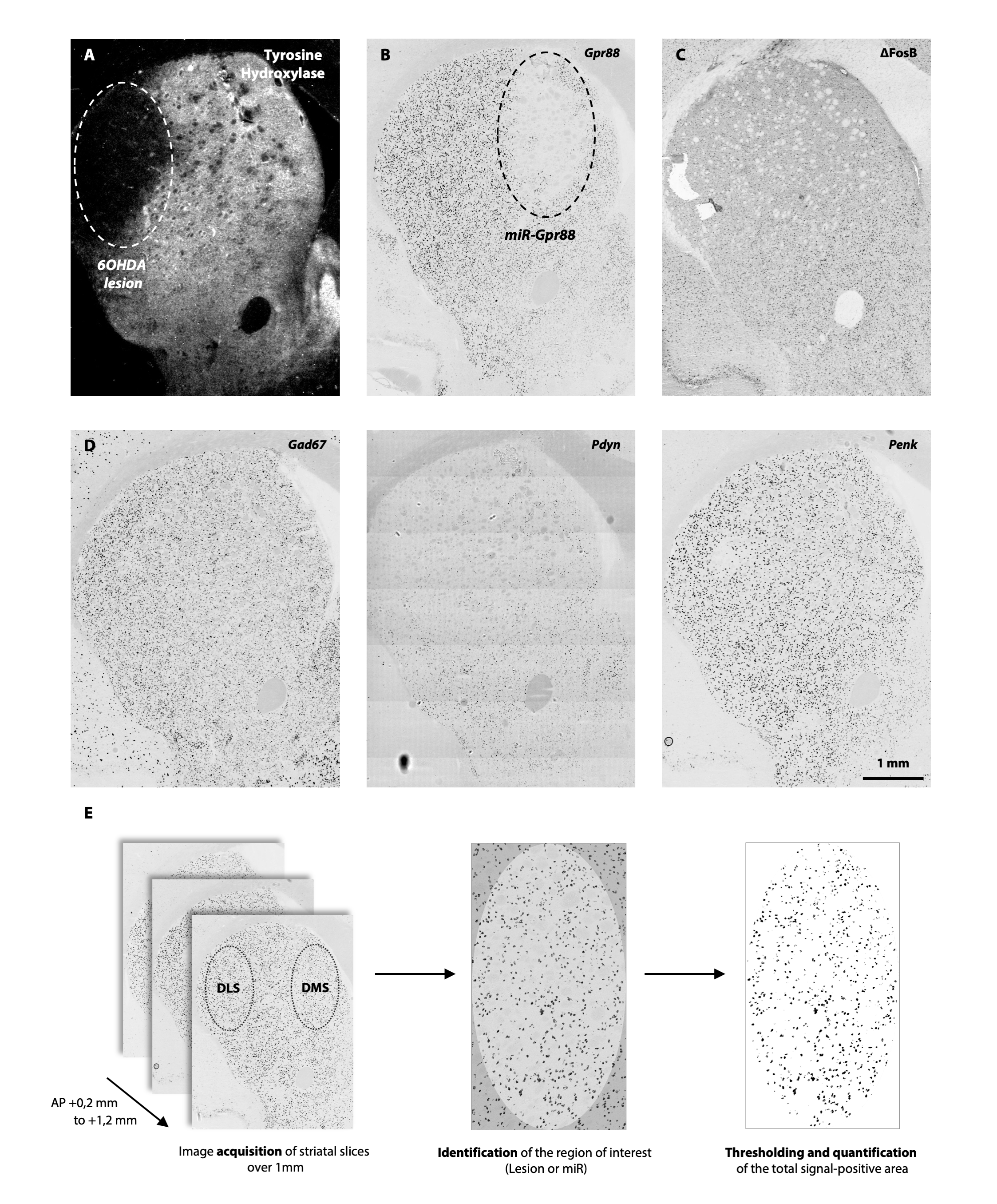

### Supplementary figure 3

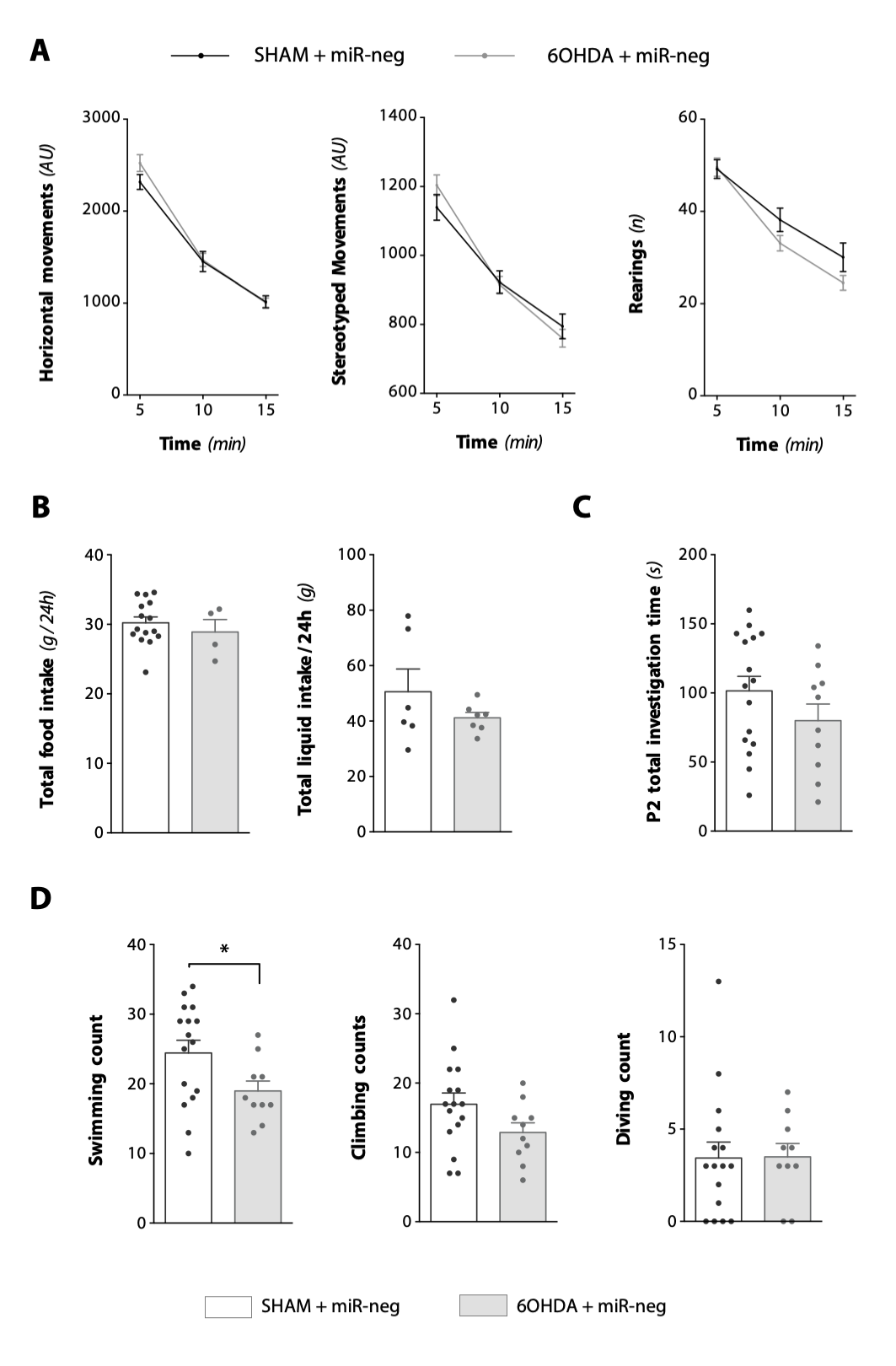

### Supplementary figure 4

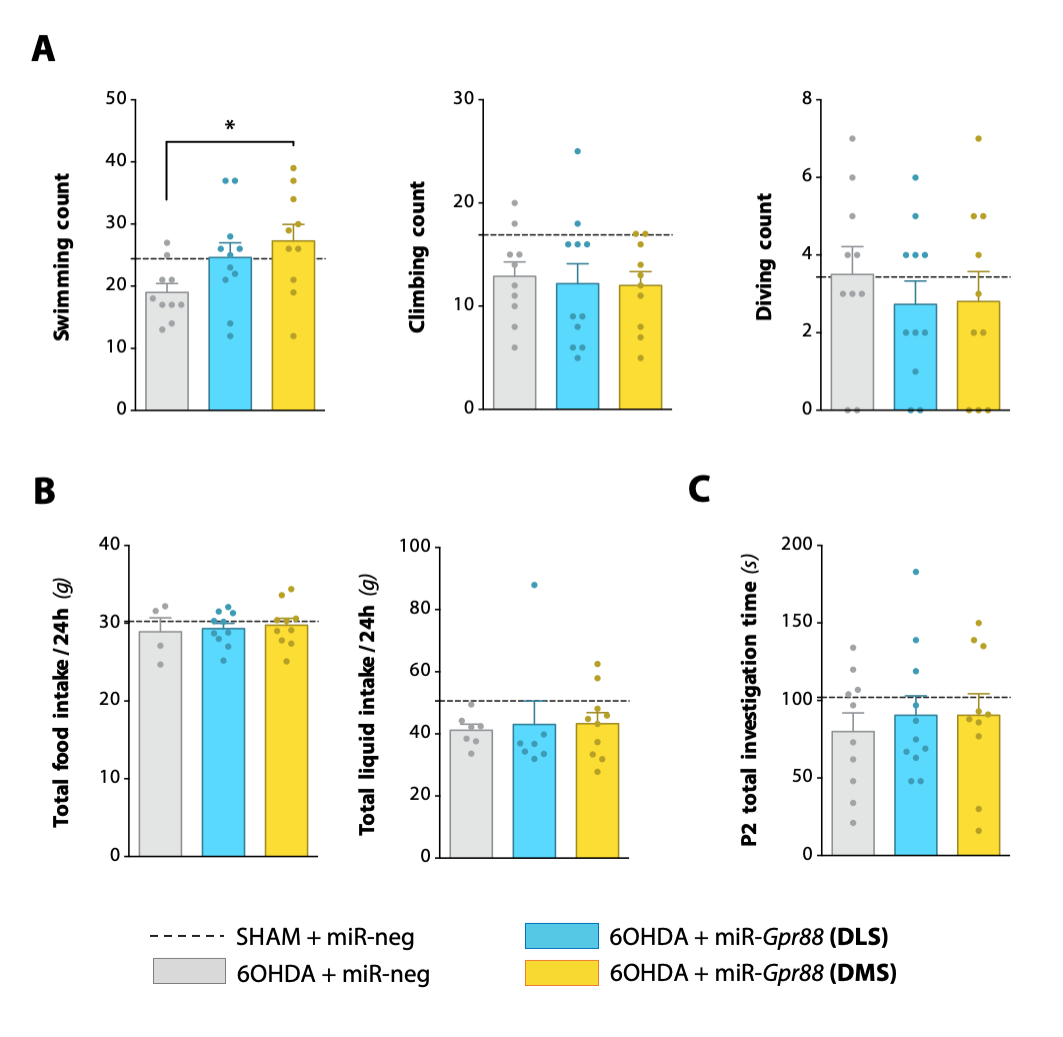

### Supplementary figure 5

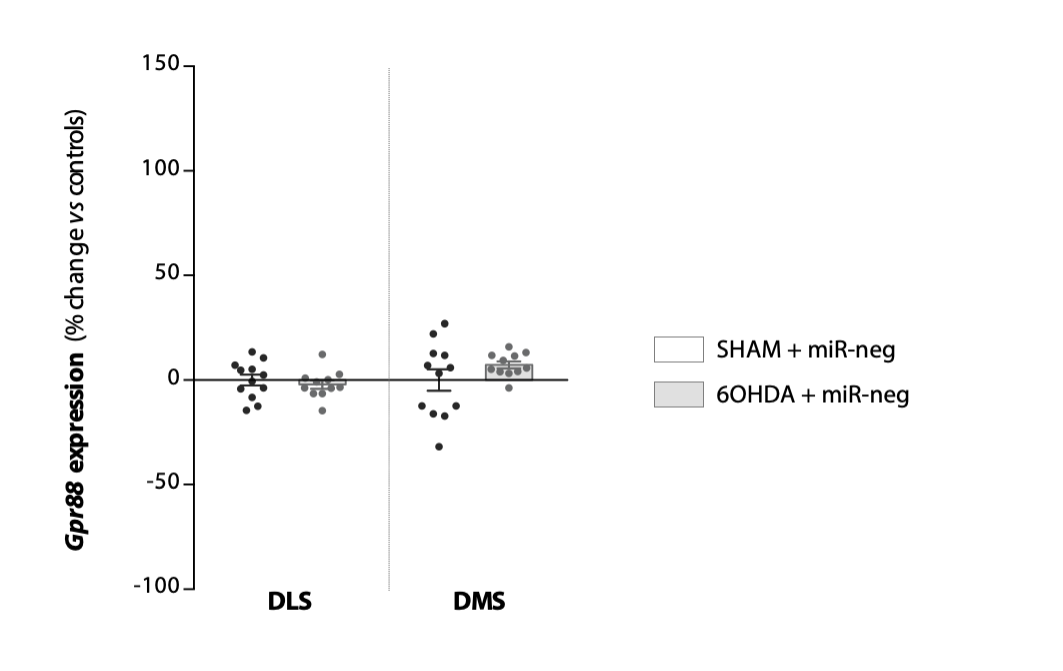
